# Tunneling nanotubes contribute to the stroma-mediated imatinib resistance of leukemic cells

**DOI:** 10.1101/425041

**Authors:** Marta D. Kolba, Wioleta Dudka, Monika Zaręba-Kozioł, Agata Kominek, Paolo Ronchi, Laura Turos, Jakub Wlodarczyk, Yannick Schwab, Dominik Cysewski, Katja Srpan, Daniel M. Davis, Katarzyna Piwocka

## Abstract

Intercellular communication within the bone marrow niche significantly influences leukemogenesis and the sensitivity of leukemic cells to therapy. Tunneling nanotubes (TNTs) are a novel mode of intercellular cross-talk. They are long, thin membranous protrusions that enable the direct transfer of various cargo between cells. Here we show that TNTs are formed between leukemic and bone marrow stromal cells. Fluorescence confocal microscopy with 3D reconstructions, correlative light-electron microscopy and electron tomography provided evidence that TNTs transfer cellular vesicles between cells. The quantitative analysis demonstrated that the stromal cells stimulate TNT-mediated vesicle transfer towards leukemic cells. Transfer of vesicular cargo from stromal cells correlated with increased resistance to anti-leukemic treatment. Moreover, specific sets of proteins with a potential role in survival and the drug response were transferred within these vesicles. Altogether, we found that TNTs are involved in the leukemia-stroma cross-talk and the stroma-mediated cytoprotection of leukemic cells. Our findings implicate TNT connections as a possible target for therapeutic interventions within the leukemia microenvironment to attenuate stroma-conferred protection.

## Introduction

Chronic myeloid leukemia (CML) is a clonal myeloproliferative disorder that accounts for 15% of all leukemias in adults. It results from a reciprocal chromosomal translocation (t[9;22][q34;q11]; Rowley, 1973) that gives rise to a bcr-abl fusion gene that is present on an abnormally short chromosome 22, known as the Philadelphia chromosome. The BCR-ABL protein is a constitutively active tyrosine kinase that is able to autophosphorylate and capable of uncontrolled signaling to numerous downstream proteins, resulting in the dysregulation of biological processes (e.g., proliferation, adherence, and apoptosis; Deininger *et al*, 2000). In the 1990s, the selective tyrosine kinase inhibitor STI 571 (imatinib) was developed to effectively block the adenosine triphosphate binding pocket of the Abl1 kinase domain, thereby inhibiting the downstream effects of uncontrolled kinase activity of the BCR-ABL protein. Since the introduction of imatinib as a first-line drug for the treatment of CML (Druker *et al*, 2001a, 2001b), the annual mortality rate has decreased from 15-20% to 2% (Alvarez *et al*, 2007). This therapy is successful in the chronic phase of the disease. However, most patients do not achieve a complete cytogenetic response and instead present residual CML disease and progression to the blast phase. The common pattern of the significant fall of the population of circulating blasts is accompanied by a reduction or delay in the decrease in the bone marrow blast population (Weisberg *et al*, 2008). Importantly, cells that are isolated and cultured *in vitro* lose their chemoresistant phenotype, suggesting a protective role of the bone marrow microenvironment (Seke Etet *et al*, 2012). Several studies have investigated this issue, demonstrating that CML survival upon imatinib treatment is mediated by factors that are secreted by mesenchymal stromal cells (Bewry *et al*, 2008; Weisberg *et al*, 2008; Kumar *et al*, 2017a) and adhesion to either the extracellular matrix (Lundell *et al*, 1996) or mesenchymal stromal cells in marrow (Zhang *et al*, 2013). Additionally, different studies, including our previous studies, have shown that stromal cells are also modified and remodeled to support leukemogenesis by factors that are released by leukemic cells, indicating bidirectional interactions (Krause & Scadden, 2015; Duarte *et al*, 2018; Podszywalow-Bartnicka *et al*, 2016). Overall, intercellular communication between CML cells and bone marrow stromal cells has been identified as an important player in the treatment resistance of CML. However, no holistic vision of different ways in which stromal cells can interact with leukemic cells as well as impact of the CML microenvironment on the disease course has been proposed to date.

Tunneling nanotubes (TNTs) were discovered in 2004 as structures that mediate intercellular organelle transfer (Onfelt *et al*, 2004; Rustom *et al*, 2004). They are long and thin membranous channels that interconnect cells over relatively long distances. In one of the first original papers in which TNT were discovered they were characterized as open membrane conduits (Rustom *et al*, 2004). Additionally synaptic nanotubes formed between immune cells were discovered and characterized (Onfelt *et al*, 2004; Sowinski *et al*, Nat Cell Biol 2008; Chauveau *et al*, 2010) - they contain a submicron scale junction which enables transfer of cargo and supports immune synapses. Cellular vesicles (Rustom *et al*, 2004; Onfelt *et al*, 2006), organelles (Vallabhaneni *et al*, 2012; Yasuda *et al*, 2011; Abounit *et al*, 2016b; Ahmad *et al*, 2014), miRNAs (Thayanithy *et al*, 2014), viral particles (Sowinski *et al*, Nat Cell Biol 2008) and proteins (Schiller *et al*, 2013; Costanzo *et al*, 2013) can be actively transported directly from cell to cell, travelling along the actin or microtubule backbone of TNTs. Soon after their discovery, TNTs were recognized as a novel mode of intercellular communication (Davis, Sowinski 2008; Ariazi *et al*, 2017; Baker, 2017; Nawaz & Fatima, 2017). They have been found in a plethora of cell types to date, including immune cells (Önfelt *et al*, 2004; Onfelt *et al*, 2006), neurons (Gousset *et al*, 2009), cardiomyocytes (Figeac *et al*, 2014), endothelial cells (Liu *et al*, 2014; Yasuda *et al*, 2010), mesenchymal stromal cells (Pasquier *et al*, 2013), and cancer cells (Lou *et al*, 2012b; Thayanithy *et al*, 2014; Pasquier *et al*, 2012). Their involvement in the pathogenesis of such diseases as neurodegenerative diseases (Costanzo *et al*, 2013; Abounit *et al*, 2016a), prion diseases (Gousset *et al*, 2009; Abounit *et al*, 2016b), viral infections (Lachambre *et al*, 2014; Kumar *et al*, 2017c) and numerous cancers (Lou *et al*, 2012b; Pasquier *et al*, 2013; Desir *et al*, 2016) has been documented. They have also been identified in leukemia. Tunneling nanotubes have been shown to be formed between acute myeloid leukemia (AML) cells (Omsland *et al*, 2017) and between AML cells and bone marrow stromal cells (Moschoi *et al*, 2016). The latter resulted in the transfer of mitochondria toward AML cells, suggesting a clear survival advantage for AML cells. Acute lymphoid leukemia (ALL) cells have also been shown to signal to mesenchymal stromal cells (MSCs) through TNTs, triggering the secretion of prosurvival cytokines (Polak *et al*, 2015). A follow-up study showed that mitochondria, transmembrane proteins, and autophagosomes can be transported from AML cells toward MSCs (de Rooij *et al*, 2017). Overall, these studies suggest that leukemic cells employ TNT-dependent signaling to modulate their microenvironment, but in-depth analyses of molecular transfer and the functional consequences of TNT-mediated intercellular transfer are lacking. Moreover, the role of TNT-mediated transfer in acquiring resistance to treatment has not been investigated in the broader context of other leukemias, including BCR-ABL-positive CML.

The present study found the presence of TNTs that were formed between CML cells and bone marrow stromal cells. We observed TNT-dependent cellular vesicle transfer from stromal cells toward CML cells, resulting in protection against imatinib-induced apoptosis. We also identified sets of proteins that were exchanged between stromal cells and CML cells within the process of intercellular protein transfer (Ahmed & Xiang, 2011; Rechavi *et al*, 2009) and sets of proteins that transferred toward CML cells specifically in a TNT-dependent manner. Biological processes that can be modulated by TNT-dependent protein transfer might confer the survival of leukemic cells and their resistance to treatment.

## Results

### Tunneling nanotubes are formed between CML cells and stromal cells

Intercellular communication within the bone marrow niche is an important element that influences leukemogenesis and the biology of leukemic cells. Previous studies demonstrated the possibility of intercellular communication through the formation of TNTs, thin membrane connections that enable direct communication between cells (Ariazi *et al*, 2017; Ady *et al*, 2014; Lou *et al*, 2012a; Gerdes *et al*, 2013). We investigated whether these connections can be formed within the leukemia microenvironment and participate in leukemia-stroma cross-talk.

We first utilized the HS-5 cell line that originates from human bone marrow stromal cells and is widely used as a model bone marrow niche cell line (Vangapandu *et al*, 2017; Weisberg *et al*, 2008). We observed the efficient formation of TNTs between HS-5 bone marrow stromal cells (Fig. 1A). The three-dimensional (3D) reconstruction of fluorescence confocal images and scanning electron microscopy (SEM) observations showed that TNTs, in contrast to filopodia, are formed between cells and did not touch the substratum (Fig. 1A, B). Next, we investigated the dynamics of TNT formation using time-lapse fluorescence confocal microscopy. Tunneling nanotubes were formed within minutes after direct cell-cell contact followed by cell dislodgement (Fig. 1C, Supplementary Movie S1). Analysis of cytoskeletal components revealed that all TNTs contained F-actin, but also microtubules were identified in some of them (Fig. 1D). The average diameter of TNTs was 0.37 ± 0.02 µm with an average length of 10-90 µm, measured by confocal microscopy in living cells. The average diameter of the TNTs was 0.21 ± 0.01 µm with an average length of 13-80 µm, measured by SEM (Supplementary Fig. S1A-D).

**Figure 1.**
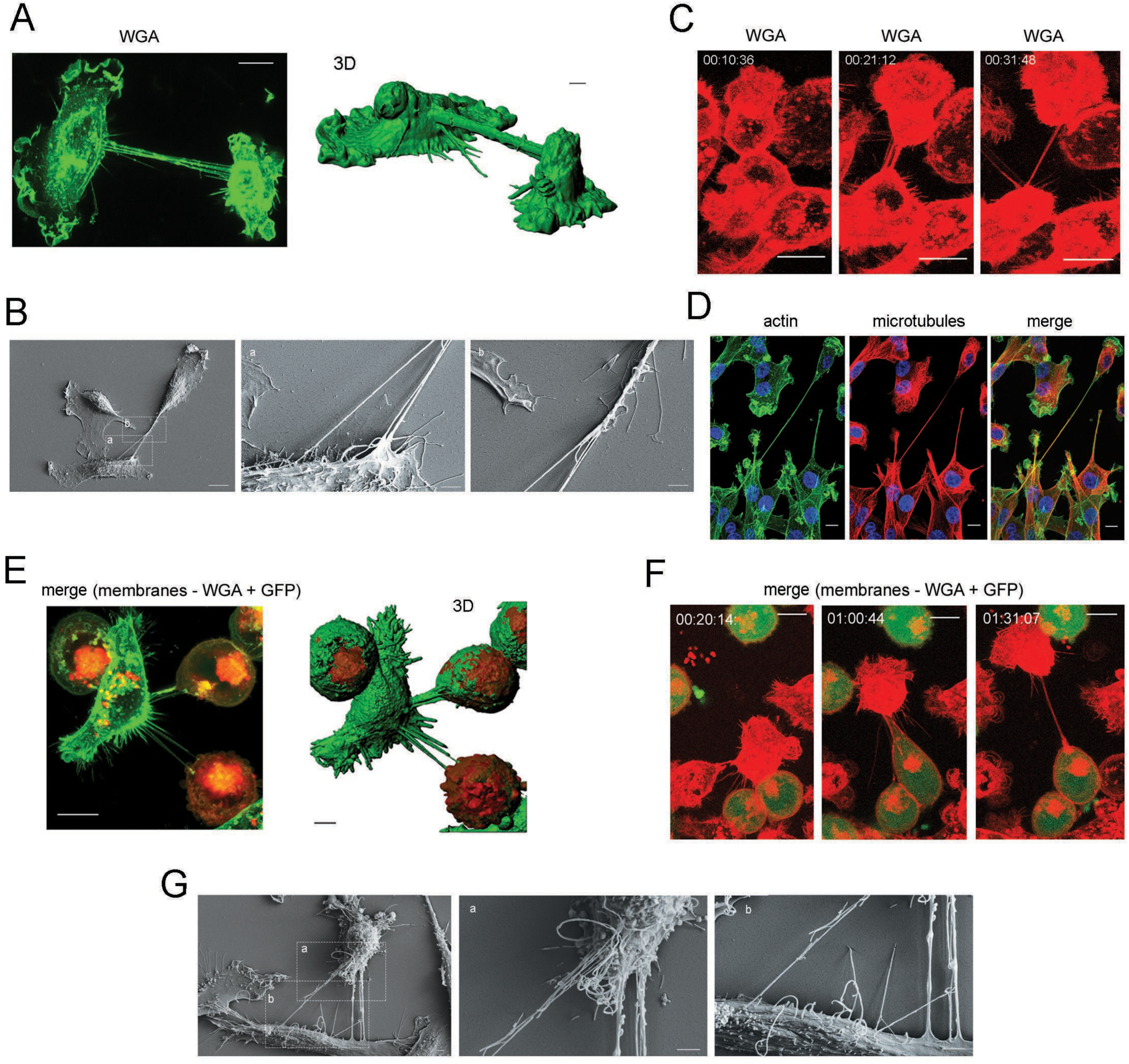
Formation of tunneling nanotubes (TNTs) between leukemic cells and stromal cells. **A.** Representative image of TNTs that interconnected stromal cells (left) and 3D reconstruction that shows that TNTs did not adhere to the substratum (right). Cell membranes were stained with WGA-Alexa Fluor 488. Scale bar = 4 μm. **B.** Scanning electron microscopy image of TNTs that formed between stromal cells. Zoomed images show a TNT end that protruded from the cell body and a bifurcation of the TNT. Scale bars: 10 μm (left image), 2 μm (middle and right images). **C**. Selected frames from a time-lapse experiment (h:min:sec) that presents TNT formation between stromal cells. Cell membranes were stained with WGA-Alexa Fluor 647. **D.** Representative image showing the actin (green) and microtubule (red) present inside a TNT that interconnected stromal cells. Blue indicates nuclei. The right panel shows the overlay. **E.** Representative image of heterotypic TNTs that interconnected one stromal cell and two CML cells. CML cells were previously labeled with DiD (red) (left). All cell membranes were stained with WGA-Alexa Fluor 488 (green) directly before imaging. The 3D reconstruction shows the lack of adhesion to the substratum of these structures (right). Scale bar = 4 μm **F.** Selected frames from a time-lapse experiment (h:min:sec) that presents TNT formation between stromal cells and CML cells. Red indicates all cell membranes that were stained with WGA-Alexa Fluor 647. Green indicates CML cells that expressed cytoplasmic GFP. Scale bars = 10 μm **G.** Scanning electron microscopy image of TNTs that formed between stromal cells and CML cells. Zoomed images shows that TNT ends protruded from cell bodies of stromal cells and CML cells. Scale bars = 2 μm.

We then evaluated whether TNTs can also be formed between stromal cells and CML cells (i.e., the K562 cell line), thus enabling direct interactions between both cell types. To investigate the formation of heterotypic TNTs, we set up a co-culture system at a 1:1 ratio of green fluorescent protein (GFP)-positive CML cells with stromal cells. The staining of membranes in living cells with WGA-Alexa Fluor 647 revealed the presence of similar structures that connected distant cells of these two types (Fig. 1E). Time-lapse confocal microscopy showed that heterotypic TNTs formed within minutes upon direct cell contact followed by cell dislodgement (Fig. 1F, Supplementary Movie S2), and they did not touch the substratum (Fig. 1E, G). They did not significantly differ in diameter or length from homotypic TNTs that were formed by stromal cells (Supplementary Fig. S1A-D).

To quantify the propensity of both cell types to form TNTs, we acquired confocal z-stack images of living cells (10 fields of view per condition) that were grown under mono- or co-culture conditions and calculated the average number of TNTs per 100 cells. We found that CML cells were much less prone to form TNTs between themselves than stromal cells. Only 1.0 ± 0.6 TNTs among 100 CML cells were observed, in comparison to 150.0 ± 32.0 TNTs that were formed by 100 stromal cells. However, upon co-culture with stromal cells, CML cells had a tendency to form more TNTs. We observed an average of 35.0 ± 6.0 homo-and heterotypic TNTs that were formed per 100 CML cells (Fig. 2A). In co-culture, heterotypic TNTs generally accounted for as much as 16.5% ± 3.5% of all TNTs that were identified, whereas homotypic TNTs that interconnected CML cells constituted only 1.8% ±1.0% of the overall number (Fig. 2B). Importantly, heterotypic TNTs that formed between CML cells and stromal cells constituted 92.1% ± 4.5% of all TNTs that were formed by CML cells. Only a fraction of the cells at a given time point were connected by TNTs, with there often being more than one nanotube per cell (≤ 5 per CML cell and ≤ 15 per CML cell; Fig. 2C).

**Figure 2.**
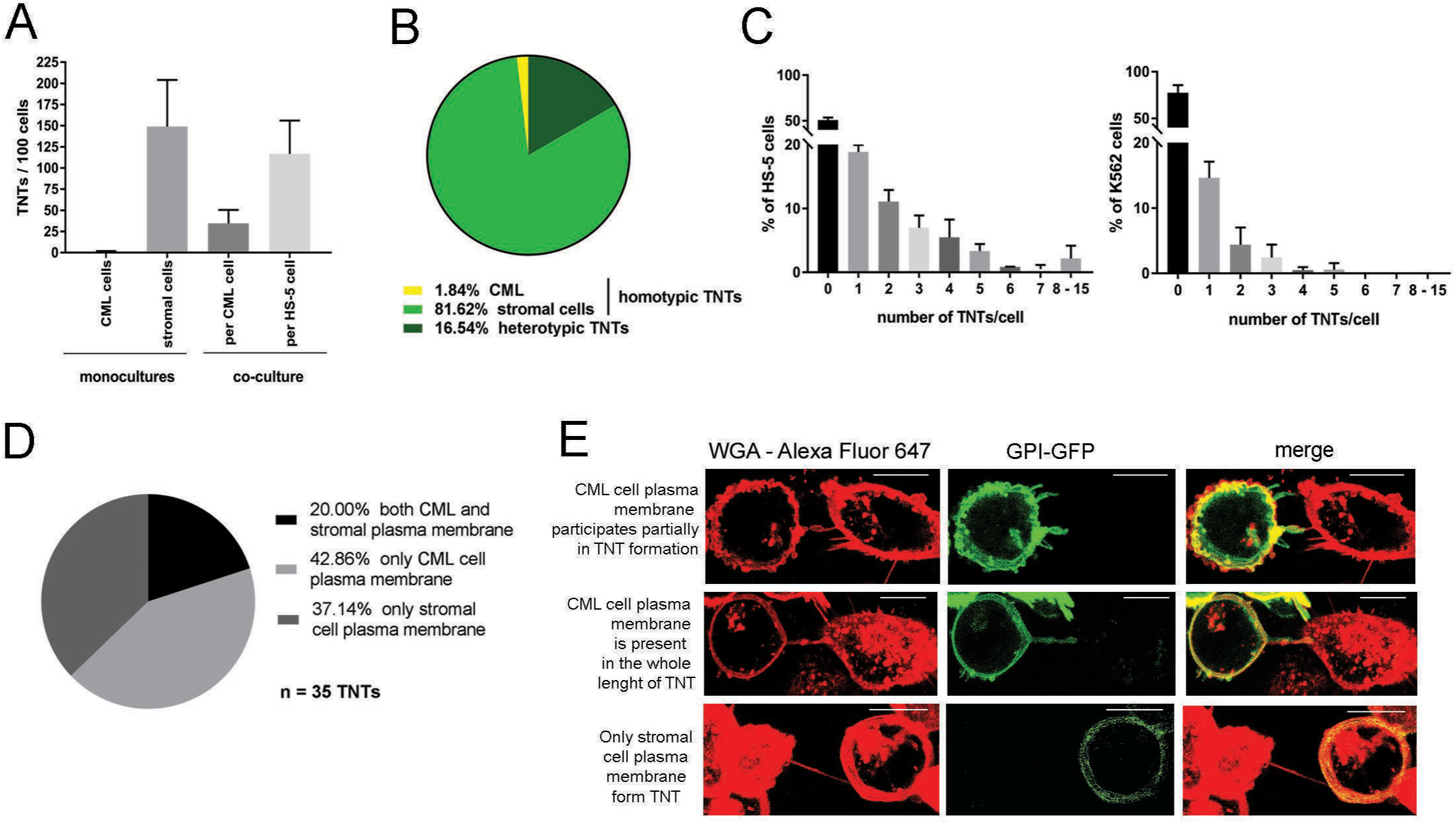
Distribution of tunneling nanotubes (TNTs) that formed in mono- and co-cultures of stromal cells and leukemic cells. **A.** Average numbers of TNTs per 100 cells quantified in mono- and co-culture set-ups by confocal microscopy after 24 h of cell culture. The graph shows that stromal cells had a propensity to form TNTs, whereas CML cells were prone to form TNTs only upon co-culture with stromal cells. **B.** Prevalence of homo- and heterotypic TNTs in a co-culture set-up, quantified by confocal microscopy. The diagram shows that the majority of TNTs that were found in co-cultures were homotypic TNTs that interconnected stromal cells. Heterotypic TNTs constituted a considerable portion of overall TNTs. CML cells in co-culture were significantly more involved in TNT formation with stromal cells than in TNT formation with other CML cells. **C.** Percentage of stromal cells (left) and CML cells (right) that exhibited a given number of TNTs in a co-culture set-up. Stromal cells and CML cells usually formed more than one TNT per cell, and stromal cells had many more TNTs per cell compared with CML cells. **D.** Participation of plasma membranes of stromal cells and CML cells in heterotypic TNT formation, quantified by confocal microscopy after 48 h of co-culture. CML cells were transfected by nucleofection with a GPI-GFP-encoding plasmid, FACS sorted, and subjected to co-culture with stromal cells. All cell membranes were stained with WGA-Alexa Fluor 647 directly before imaging. **E.** Representative images of heterotypic TNTs that were formed by plasma membranes of both stromal cells and CML cells (upper panel) or exclusively by plasma membranes of CML cells (middle panel) or stromal cells (lower panel). Red indicates WGA-Alexa Fluor 647 in CML cells. Green indicates GPI-GFP in CML cells. Scale bars = 10 μm. All of the data are expressed as the mean ± SEM of three independent experiments.

We found that leukemia cells had a lower ability to form TNTs. We investigated whether both cell types or only one cell type participates in the formation of heterotypic TNTs. Therefore, CML cells were transfected with a plasmid that encoded GFP with a glycosylphosphatidylinositol (GPI) tag, and TNT formation was assessed by confocal microscopy. All plasma membranes were stained with WGA-Alexa Fluor 647. Thus, the membranes of both cell types showed red fluorescence however only leukemia cells’ membrane showed green GFP fluorescence, enabling us to estimate the involvement of leukemia’s component. We found that in 20% of heterotypic TNTs, membranes of both CML cells and stromal cells participated in TNT formation (Fig. 2D, E). Additionally, in heterotypic TNTs formed between CML and stromal cells, the CML membranes were found in the whole length of 43% counted TNTs or did not participated in the TNT formation in 37% of them, respectively. The length of TNTs did not depend on the origin of the plasma membrane (Supplementary Fig. S2). These results clearly showed that CML cells were involved in heterotypic TNT formation equally to stromal cells.

Altogether, the above data revealed that TNTs were formed between stromal cells (homotypic) and between stromal cells and CML cells (heterotypic), and heterotypic nanotubes can be formed by membrane from either cell or both cells. However, leukemic cells became more prone to TNT formation upon co-culture with stromal cells. Despite of different ability/propensity to form TNT, both cell types actively contributed to TNT formation (Fig. 2E).

### Tunneling nanotubes mediate cargo transfer between stromal cells and CML cells

Tunneling nanotubes have been previously reported to mediate the intercellular trafficking of cellular cargo (Rustom *et al*, 2004; Burtey *et al*, 2015; Schiller *et al*, 2013; Lou *et al*, 2012b; Gurke *et al*, 2008). To evaluate possible cargo transfer between CML cells and stromal cells, we first verified the actin penetration upon active process of TNTs formation. This was assessed in co-cultures of stromal cells and CML cells that were transfected with a plasmid that enabled the visualization of F-actin in living cells. The plasma membranes of all of the cells were stained with WGA-Alexa Fluor 647. All of the observed heterotypic TNTs (*n* = 39) were partially or completely penetrated by actin in CML cells (Fig. 3A), and none of them were closed or inaccessible for actin polymerization.

**Figure 3.**
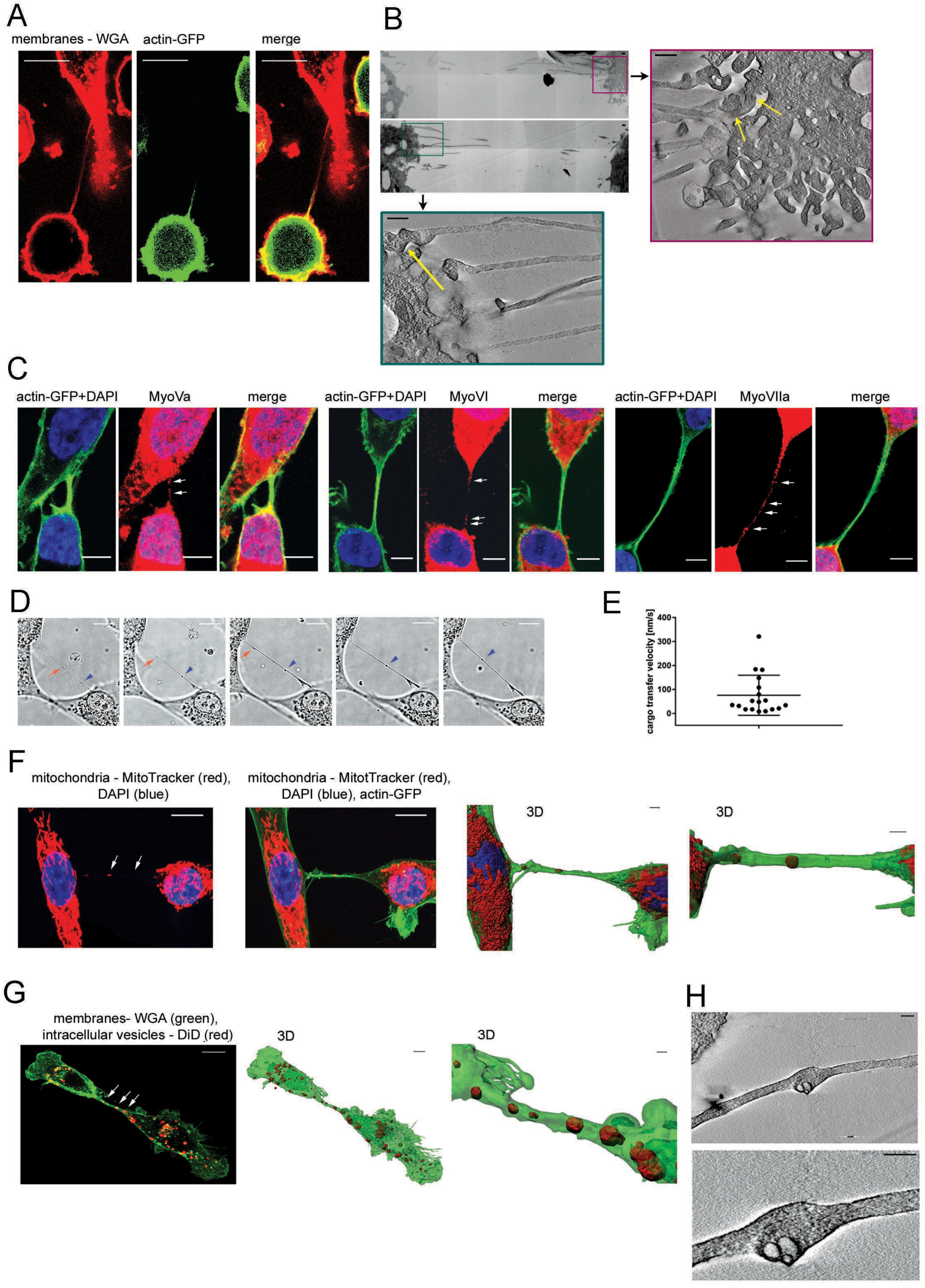
Cargo transfer mediated by tunneling nanotubes (TNTs). **A.** Representative image that shows the penetration of actin of CML cells into a heterotypic TNT after 48 h of co-culture. CML cells were transfected by nucleofection with a LifeAct-GFP-encoding plasmid, FACS sorted, and subjected to co-culture with stromal cells. All cell membranes were stained with WGA-Alexa Fluor 647 directly before imaging. Single channels and a merged image are shown. **B.** Transmission electron microscopy image that shows that TNTs provide continuity with the cell body. Scale bar = 500 nm. **C** Representative images that show the presence of molecular motors inside TNTs. Green indicates the actin cytoskeleton. Blue indicates nuclei. Red indicates antibody-stained myosin Va (left), myosin VI (middle), and myosin VIIa (right). Scale bar = 5 μm. Single channels and a merged image are shown. **D.** Frames from a time-lapse experiment that present the transport of two bulges along a TNT. **E.** Velocity of cargo transfer in the time-lapse experiments (*n* = 18 events). **F.** Representative images that show the presence of mitochondria inside TNTs. Green indicates actin. Blue indicates nuclei. Red indicates mitochondria that were stained with MitoTracker Deep Red. The 3D reconstruction shows in detail mitochondria that were present within the lumen of the TNT. Scale bar **=** 4 μm. **G.** Representative image that shows the presence of cellular vesicles inside TNTs. Green indicates cell membranes that were stained with WGA-Alexa Fluor 488. Red indicates cytoplasmic vesicles that were stained with DiD. The 3D reconstruction shows in detail cytoplasmic vesicles that were present within the lumen of the TNT. Scale bar **=** 4 μm **H.** Electron tomography images from CLEM experiment show cellular vesicles that were present inside a TNT. Scale bar = 500 nm (left image) and 200 nM (right image). Scale bars for all confocal images (except C) = 10 μm.

In order to characterize the ultrastructure of the TNTs emerging from stroma cells, we used EM tomography. As shown in Fig. 3B, the plasma membrane at the emerging site is highly convoluted, with several invaginations and protrusion of different length. Interestingly, the thickness of the emerging TNTs, which are continuous with the plasma membrane of the cell body, is remarkably consistent (~ 150 nm).

Moreover, we observed the presence of molecular motor proteins that are required for cargo trafficking. Myosins MyoVa (Rustom *et al*, 2004), MyoVI, and MyoVIIa were identified inside TNTs (Fig. 3C). It is to mention that in order to visualize myosins within the thin TNT the signal from red channel had to be increased, resulting in overexposition of cellular signal. Nevertheless, we can not exclude nuclear, in addition to cytoplasmic, localization of myosines (Karolczak *et al*, 2004; Caridi *et al*, 2018). We found that TNTs that were stained with WGA often contained bulges (Fig. 1A), indicating the movement of cargo (Onfelt *et al*, 2006; Veranic *et al*, 2008). In living cells, we detected the movement of these bulges along the TNTs (Fig. 3D, Supplementary Movie S3) and quantified their velocity (Fig. 3E). Transfer velocities remained within the range of velocities that were reported for myosin-driven movement (Pierobon *et al*, 2009). Altogether, these data strongly suggest that TNT-mediated cargo transfer is possible between CML cells and stromal cells.

The specific fluorescent tracking of organelles, followed by confocal microscopy and 3D reconstruction, allowed us to identify mitochondria (Fig. 3F) and cytoplasmic vesicles (Fig. 3G) inside TNTs. MitoTracker-stained mitochondria and DiD-stained cellular vesicles were detected and visualized upon 3D image reconstruction in the lumen of TNTs that directly connected donor and acceptor cells. These data were supported by correlative light electron microscopy (CLEM), in which we labeled plasma membranes with WGA and identified, localized on the grid, and imaged TNTs that contained bulges by confocal microscopy. We then processed the samples for TEM, relocalized the same cells, and performed 3D electron tomography imaging. The images showed the presence of vesicles within the lumen of the TNTs (Fig. 3H). These vesicles had an average diameter of 111 ± 33 nm that corresponded to the typical size of cellular vesicles.

### Cellular vesicles are bidirectionally transported between stromal cells and CML cells

The above data provide evidence that vesicles are present inside TNTs and can move within nanotubes. To quantify the active cross-talk *via* TNTs and the dynamics of the cell-to-cell transport of cellular vesicles, we used a method based on flow cytometry to analyze the transfer of fluorescently labeled vesicles between GFP-positive CML cells and stromal cells in co-culture (Fig. 4A). Such a strategy has been used before to quantify the nanotube-mediated vesicles transfer. Vesicles of donor cells were labeled with DiD, and the acceptor cell population remained unstained. DiD stains lipophilic structures in the cell, has very low toxicity, does not undergo passive transport, and is used to track cellular vesicles that are transferred between cells by different ways, including TNTs (Honig & Hume, 1986; Daubeuf *et al*, 2009; Onfelt *et al*, 2006; Gurke *et al*, 2008). To exclude the possibility of indirect extracellular transfer of vesicles between cells, control experiments were performed in which acceptor cells were either treated with conditioned medium from donor cells or were physically separated from donor cells by a filter with 1.0 µM pores that enabled the cells to share the same media but excluded the possibility of physical direct contact. The transfer of cellular vesicles from donor to acceptor cells was measured as a percentage of acceptor cells that gained fluorescence, corresponding to vesicles after 0, 6, 24, and 30 h of co-culture with donor cells. An analogous experiment was performed with monocultures of CML and stromal cells. The percentage of DiD+ cells, representing acceptor cells that obtained fluorescent cargo, was estimated. An example flow cytometry analysis is presented in Fig. 4A. We observed the exchange of cellular vesicles that occurred in all of the experimental set-ups. The transfer efficiency depended on both the donor and acceptor cell types (Fig. 4B). In mono-cultures, stromal cells were efficient donors (i.e., 75.0% ± 3.3% of acceptor cells received fluorescent vesicles), whereas CML cells were poor donors when CML cells also served as acceptors (i.e., 9.0% ± 0.5% of acceptor cells received fluorescent vesicles). In co-culture, we observed the bidirectional exchange of cargo with different efficiencies, depending on the direction of transfer (20% ± 1% of stromal cells *vs*. 51.0% ± 1.5% of CML cells received fluorescent vesicles after 24 h of co-culture). CML cells were significantly more efficient in trafficking vesicles toward stromal cells (20% ± 1%) than toward other CML cells (9.0% ± 0.5%). These data clearly correlated with the counts of homo- and heterotypic TNTs that were formed by CML and stromal cells in mono- and co-cultures (compare Fig. 4C with Fig. 2A). Such a correlation, together with the necessity for direct cell-cell contact, indicated that TNTs might be responsible for the trafficking of cellular vesicles.

**Figure 4.**
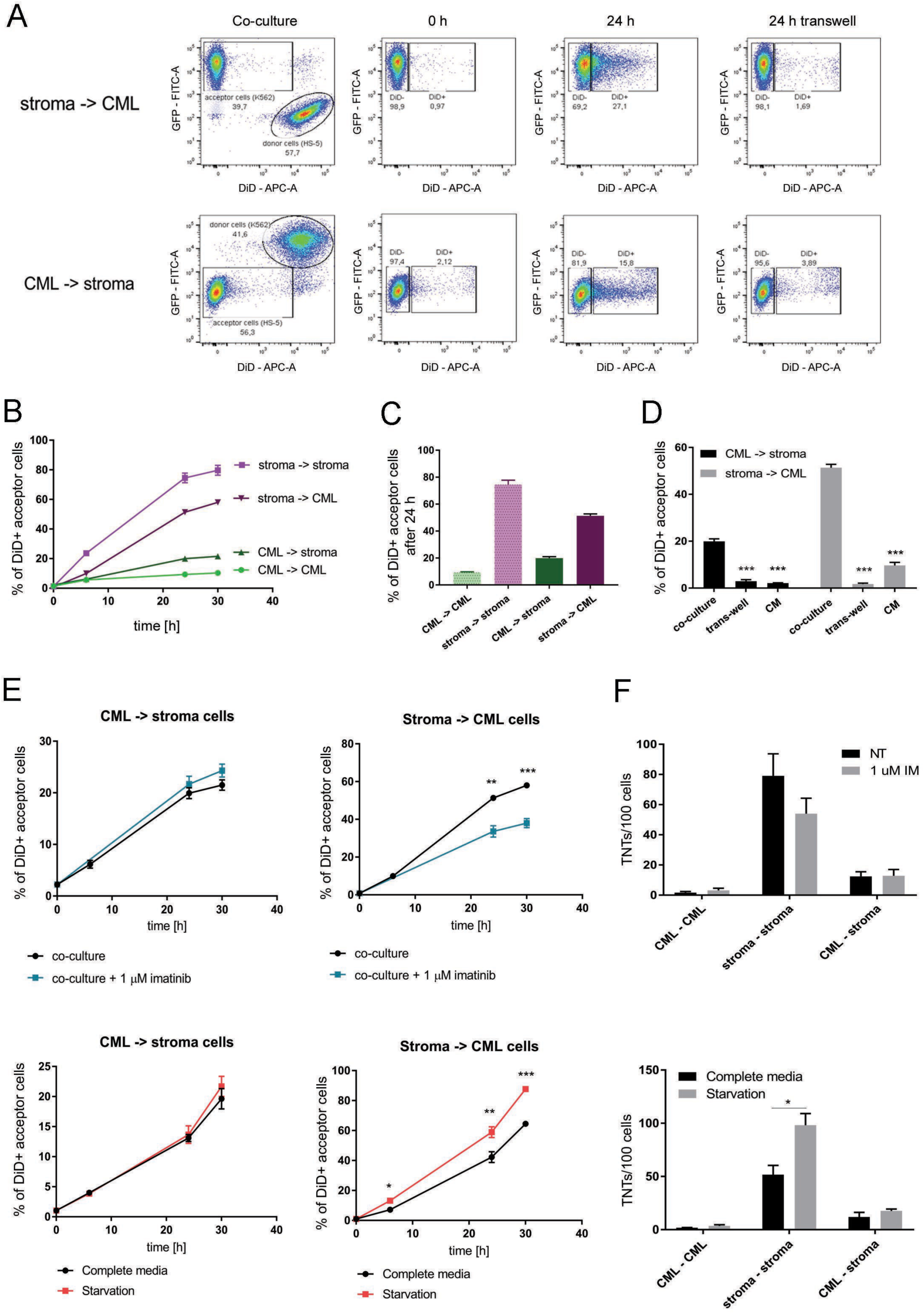
Quantitative analysis of the direct exchange of cellular vesicles between stromal cells and leukemic cells. **A.** Dot plots that present the gating strategy for the flow cytometry experiments on the exchange of cellular vesicles in a co-culture set-up. CML cells expressed cytoplasmic GFP. Donor cells were labeled with DiD for cytoplasmic vesicles. The plots depict the shift in fluorescence in acceptor cells that was caused by the uptake of fluorescently labeled vesicles. The transwell system was used as a control to show that vesicles transfer was directly contact-dependent. **B.** Time course of cellular vesicle trafficking in the mono- and co-culture set-ups, quantified by flow cytometry. The percentage of DiD+ acceptor cells is shown. **C.** Efficiency of cellular vesicle trafficking in the mono- and co-culture set-ups, quantified by flow cytometry and presented as the average percentage of acceptor cells that received fluorescently labeled vesicles that were transferred from donor cells after 24 h of culture. **D.** Direct contact-dependence of vesicle transfer. For the control experiments, (*i*) a transwell system was used to physically separate donor and acceptor cells that shared the same medium, or (*ii*) acceptor cells were cultured in conditioned medium (CM) that was collected from donor cells. **E, F.** Influence of starvation-mimicking conditions (upper row) and imatinib treatment (lower row) on (**E**) the efficiency of vesicle transfer between stromal cells and CML cells and (**F**) TNT formation in a co-culture set-up. Both treatments regulated only one direction of the bidirectional exchange of vesicles that occurred through heterotypic TNTs, which was not accompanied by a change in the prevalence of heterotypic TNTs in co-culture. All of the data are expressed as the mean ± SEM of three independent experiments.

To directly assess and exclude the role of secretion and extracellular vesicles in the vesicle transfer that was observed, we cultured acceptor cells in the 24 h conditioned medium (CM) of donor cells that were labeled for cellular vesicles and thus contained all types of secreted vesicles (Fig. 4D, CM). After 24 h, the fluorescence of vesicles that was measured in acceptor cells that were treated with CM was substantially lower than the transfer that occurred under co-culture conditions, thus allowing direct cell-cell contact (Fig. 4D). Only 2.1% ± 0.2% of stromal cells acquired fluorescent vesicles from CML cells compared with 20.0% ± 1.1% in the classic co-culture. In the reverse set-up, only 9.7% ± 1.3% of CML cells received vesicles from stromal cells, whereas this percentage reached 51.4% ± 1.5% in co-culture. Vesicle transfer in the transwell system was significantly lower compared with the regular co-culture model (2.9% ± 0.7% *vs*. 20.0% ± 1.1% for CML-to-stroma transfer and 1.7% ± 0.4% *vs*. 51.4% ± 1.5% for stroma-to-CML transfer; Fig. 4D, transwell). Altogether, these data show that the stroma-leukemia intercellular vesicle trafficking is direct, cell-to-cell contact-dependent and acts independently of intercellular signaling *via* secreted extracellular vesicles.

To further investigate the possible regulatory mechanisms that govern vesicle transfer, we subjected co-cultures to different stressful stimuli. Starvation was previously reported to stimulate formation of TNTs (Lou *et al*, 2012b). The other stimulus, imatinib, was specifically directed toward CML cells, as a specific inhibitor of BCR-ABL kinase and a first-line drug for CML treatment. We analyzed vesicle transfer by flow cytometry at different time points. Both stressful stimuli exerted an effect that was limited to only one direction of transfer (i.e., when stromal cells were donors and when CML cells were acceptors; Fig. 4E). Interestingly, these two stressful stimuli exerted an opposite influence on the exchange from stromal cells to CML cells. Imatinib treatment downregulated vesicle trafficking 1.5-fold (33.7% ± 3.0% *vs*. 51.4% ± 1.5%), whereas starvation conditions upregulated vesicle trafficking 1.4-fold (59.0% ± 3.6% *vs*. 42.0 ± 3.6%). To determine whether this change was attributable to transfer efficiency rather than to regulation of the prevalence of TNTs, we quantified homo- and heterotypic connections in co-cultures upon imatinib treatment and starvation. The number of heterotypic TNTs was unaffected by any of these conditions (Fig. 4F).

These data showed the bidirectional exchange of cellular vesicles between CML cells and stromal cells, which was directly cell-to-cell contact-dependent and might be mediated by TNTs. Both directions of transfer differed in efficiency and the degree to which they were regulated by stressful stimuli. Therefore, they may constitute two separate modes of intercellular cross-talk. The transfer of vesicles from stromal cells to CML cells was specifically regulated by imatinib, indicating existence of the mechanism at least partially dependent on the BCR-ABL signaling.

### Protection of CML cells from imatinib-induced apoptosis by stromal cells depends on the transport of cellular vesicles

CML cells have been previously shown to be protected from imatinib-driven apoptosis upon co-culture with stromal cells (Zhang *et al*, 2013; Kumar *et al*, 2017b). The protective effect of stroma was also confirmed in our experimental model (Supplementary Fig. S3). Thus, we investigated whether the TNT-mediated transfer of vesicles confers this protection. Stromal cells were labeled with DiD for cellular vesicles and co-cultured with CML cells with increasing doses of imatinib. Flow cytometry gating was used to analyze the percentage of apoptotic cells in two groups separately: (*i*) CML acceptor cells that absorbed the fluorescently labeled vesicles (*DiD+*) from donor stromal cells and (*ii*) acceptor CML cells that did not received fluorescent vesicles (*DiD-*). We found that the uptake of vesicles from stromal cells correlated with increased protection against imatinib-driven apoptosis. Namely, among CML cells that were positive for donor-derived vesicles (*DiD+*; Fig. 5A), we observed significantly fewer cells that expressed apoptotic marker (Annexin V) compared with the population of CML cells that were negative for those vesicles (*DiD-*; Fig. 5A): 15.8% ± 0.9% *vs*. 26.6% ± 0.5% upon treatment with 1 µM imatinib, respectively.

**Figure 5.**
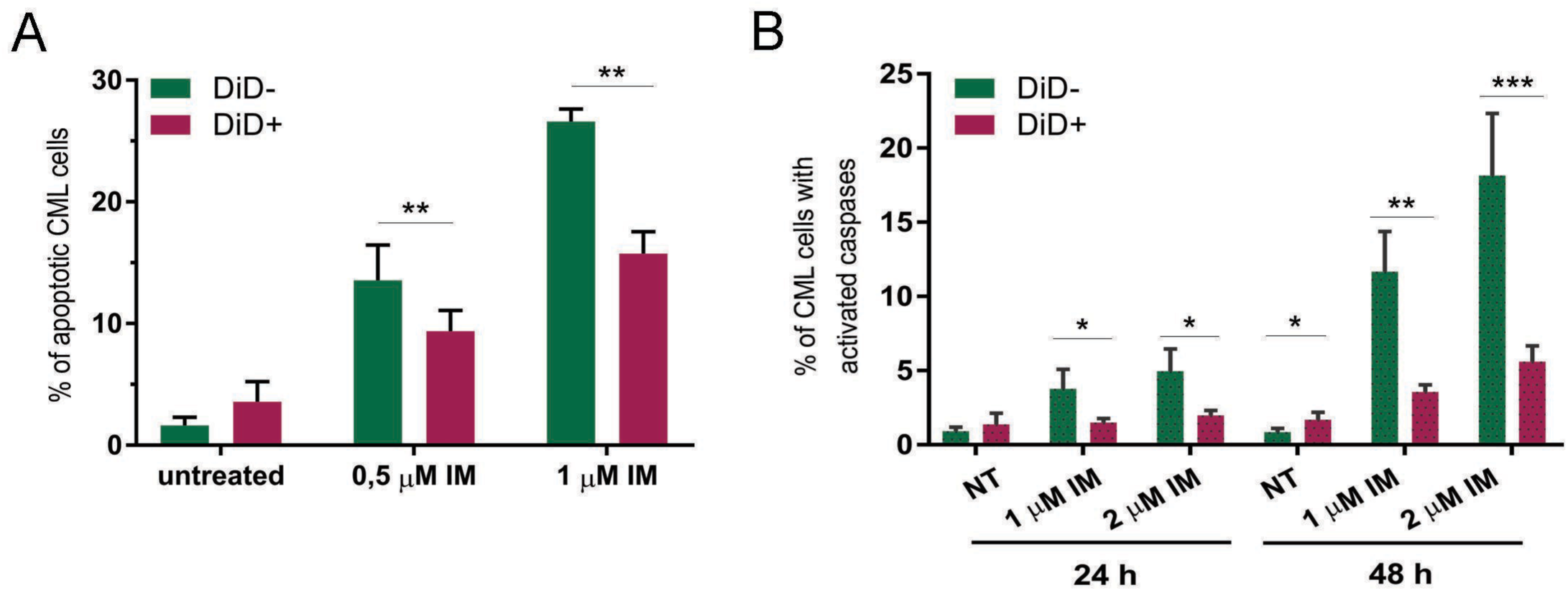
Protection of leukemic cells from imatinib-induced apoptosis by the transported vesicles. **A.** Percentage of apoptotic leukemic cells upon treatment with imatinib, analyzed separately for cells that received (DiD+) or did not receive (DiD-) cellular vesicles from stromal cells. **B.** Percentage of leukemic cells with active caspases upon treatment with imatinib, analyzed separately for cells that received (DiD+) or did not receive (DiD-) cellular vesicles from stromal cells. All of the data are expressed as the mean ± SEM of three independent experiments. Student’s *t*-test with Welch correction was used to test differences between two conditions. \**p* < 0.05, \*\**p* < 0.01, \*\*\**p* < 0.001.

Caspases are critical mediators of cell death. To support previous data, we further investigated whether the resistance to imatinib that is conferred by the direct transfer of cellular vesicles from stromal cells to CML cells would result in the modulation of general caspase activity. We performed an experiment that was similar to the experiment that is described above using a multi-caspase test to stain for active caspases-1, -2, -3, -6, -8, -9, and -10 with a single probe. The percentage of cells with active caspases was estimated after the discrimination of living cells. Caspase activity was upregulated upon imatinib treatment in CML acceptor cells; importantly, it was significantly higher in the population of CML cells that did not receive cellular vesicles from stromal cells (*DiD-;* Fig. 5B) compared with CML cells that received those vesicles (*DiD+*; Fig. 5B): 11.7% ± 1.2% *vs*. 3.6% ± 0.2% after 48 h of 1 µM imatinib treatment. These data confirmed that the direct uptake of vesicles from stromal cells correlated with increased resistance to the imatinib-induced apoptosis of leukemic cells.

### Specific sets of proteins are transported bidirectionally between stromal cells and CML cells

We found that the intercellular vesicle trafficking is directly cell-to-cell contact-dependent and acts independently of intercellular signaling *via* secreted extracellular vesicles. Thus, we hypothesized that the stroma-dependent nanotube-mediated cytoprotective effect on leukemic cells might result from proteomic exchange that occurs together with vesicle transfer.

To identify proteins that are shuttled between CML cells and stromal cells, we applied a mass spectrometry (MS)-based trans-SILAC approach (Supplementary Fig. S4). Donor cells were cultured in media that contained heavy isotopologues of lysine and arginine to allow for > 98.5% incorporation of these amino acids into their proteome (Supplementary Table S1). The cells were then co-cultured with acceptor cells that were or were not treated with imatinib. After 24 h, the GFP-positive CML cells were sorted by flow cytometry. Liquid chromatography-dual MS (LC-MS/MS) enabled to identify heavy-labeled proteins within the proteome of acceptor cells. These proteins were synthesized in donor cells and transferred from donor cells to acceptor cells during co-culture.

As a proof of concept, we performed pilot experiments to reveal the existence of specific sets of proteins that were exchanged between cells. We identified 11 proteins that were transferred from CML cells to stromal cells and 33 proteins that were transferred from stromal cells to CML cells by applying a very strict false discovery rate (FDR ≤ 0.01; Supplementary Table S2). Both sets of proteins (i.e., transferred from CML cells to stromal cells and vice versa) were grouped into functional networks of interacting proteins with very high statistical values (Supplementary Fig. S5A, B; PPI enrichment *p* = 3.77e-06 and < 1.0e-16, respectively). The GeneOntology statistical overrepresentation test showed the enrichment of several biological processes for both sets of proteins (Supplementary Fig. S5C, D). These data demonstrate the general applicability of the trans-SILAC method to identify proteins that were transferred between cells in co-culture. Initial bioinformatics analysis showed that the proteins that were identified were not randomly transported but rather comprised specific sets of proteins with a possible specific function in acceptor cells.

As we observed that the protein exchange is possible between leukemic and stromal cells, we proceeded to the main part of the study in which we applied the trans-SILAC approach to identify proteins that are present within the stroma-derived vesicles, mediate cytoprotection, and are transferred from stromal cells to CML cells. To capture in depth the set of proteins that were transferred, additional fractionation steps for MS analysis were applied. We cultured stromal cells in SILAC media as described above and stained them for cytoplasmic vesicles with DiD dye (Fig. 6). We set up the co-culture with CML cells for 24 h, followed by the fluorescence-based flow cytometry sorting of CML cells that were positive (*DiD+*) and negative (*DiD-*) for stroma-derived vesicles that were labeled with DiD.

**Figure 6.**
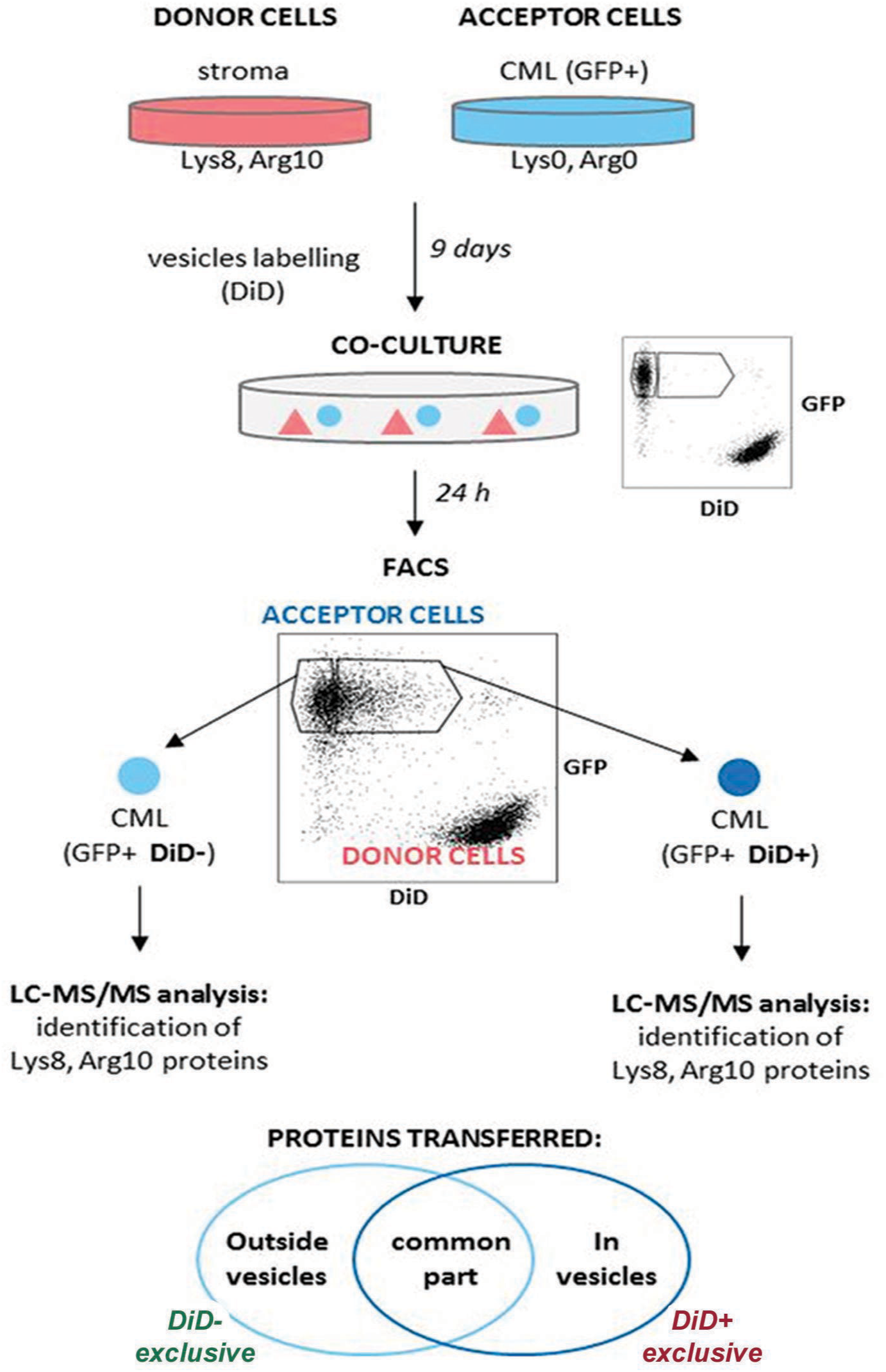
The scheme showing the mass spectrometry (MS)-based trans-SILAC approach performed to identify proteins that are present within stroma-derived vesicles. Donor stromal cells were cultured in media containing heavy isotopologues of Lys and Arg for 9 days to allow complete incorporation of these amino acids into their proteome. They were stained with DiD for cellular vesicles and subjected to co-culture with acceptor CML cells for 24 h. Afterwards two subpopulations of CML cells were sorted using FACS: cells which received (DiD+) or did not receive (DiD-) cellular vesicles from donor stromal cells. In both samples, heavy proteins, transferred from donor stromal cells, were identified. Only proteins present exclusively in the DiD+ sample were considered to have been transferred along with the studied vesicles.

We identified 481 donor-derived heavy-labeled proteins in the CML *DiD-* population and 646 in the CML *DiD+* population (Supplementary Table S2). There were 351 proteins in common between these two groups (Fig. 7A). We selected 294 proteins (named *DiD+ exclusive*) that were specifically and exclusively transferred within the studied vesicles. The remainder of the proteins were transferred either only by other means (*DiD- exclusive*) or both ways (*common*). All three protein groups had a similar molecular weight distribution (Fig. 7B), demonstrating that regardless of the transport mechanism, there was no molecular mass weight restriction.

**Figure 7.**
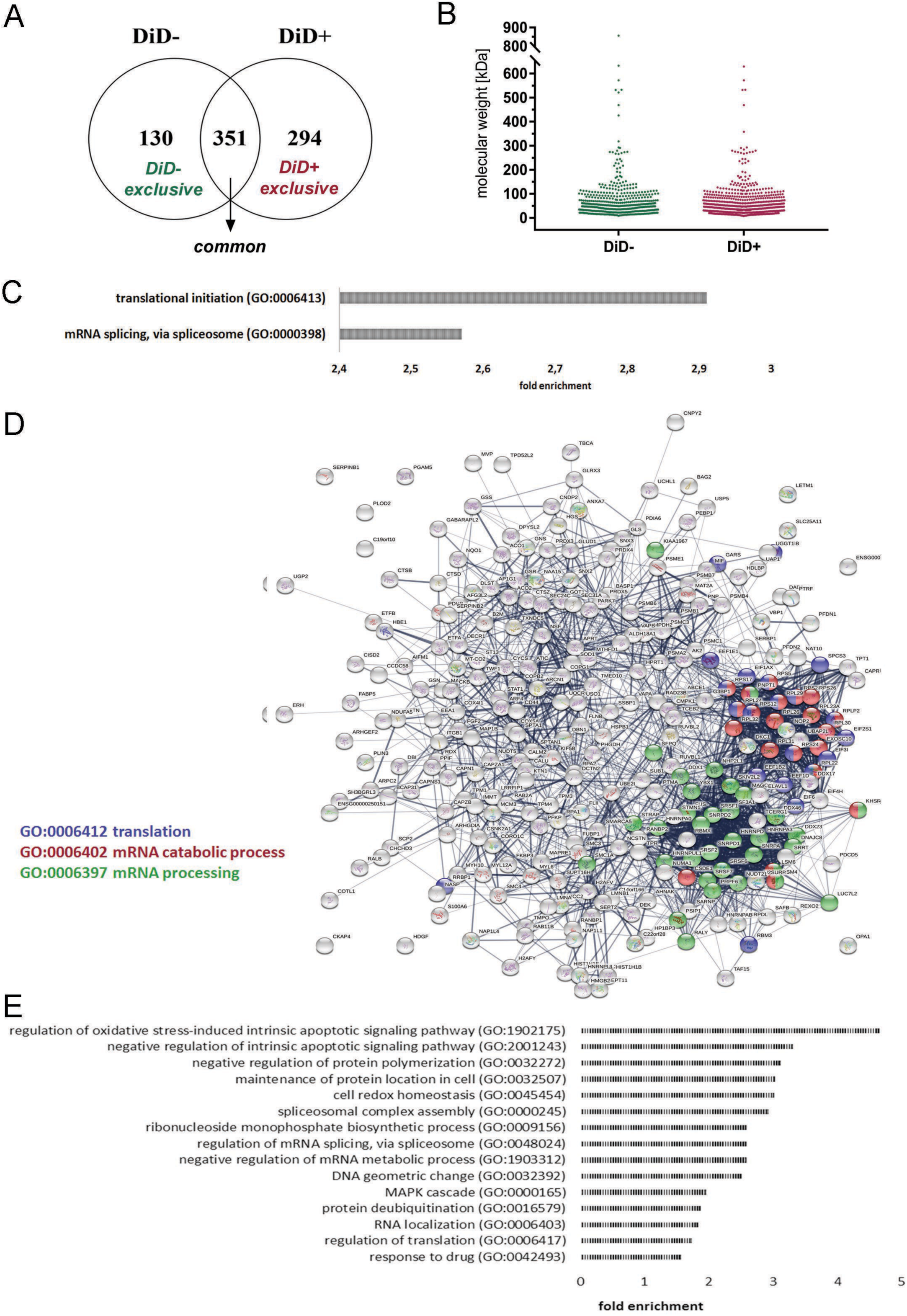
Proteins that were transported along with cellular vesicles from stromal cells to leukemic cells. **A.** Venn diagram comparison of sets of proteins that were transferred from stromal cells to CML cells either separately (DiD-) or together with DiD-labeled cellular vesicles (DiD+). **B.** Molecular weights of proteins that were transferred from stromal cells to CML cells separately (DiD-) or together with DiD-labeled cellular vesicles (DiD+). **C.** Results of statistical overrepresentation test of the *DiD+ exclusive* list of proteins (Panther GeneOntology). The list was tested for the enrichment of proteins that are involved in GeneOntology Biological Processes against the reference list of genes of the K562 cell line from the PRIDE database, completed with the list of genes that were identified in the present study. Fisher’s Exact with FDR multiple test correction was used. Only results with FDR ≤ 0.05 are presented. **D.** Confidence view of functional protein association networks (String software) of the *DiD+ exclusive* list of proteins. Hubs represent proteins. Edges represent protein-protein associations. The line thickness indicates the strength of data support. PPI enrichment *p* < 1.0e-16. **E.** GeneOntology Biological Processes that were enriched exclusively by the list of proteins that were transferred along with DiD-labeled cellular vesicles (DiD+).

The identified proteins underwent a statistical overrepresentation test using the Panther program against the reference set. Within the *DiD- exclusive* group of proteins, no GeneOntology biological processes were enriched and within the group of *common* proteins, 56 GeneOntology biological processes were enriched (FDR ≤ 0.05; Supplementary Table S3). In the *DiD+ exclusive* protein group, two biological processes were specifically enriched (Fig. 7C): translation initiation (GO: 0006413) and mRNA splicing *via* spliceosome (GO: 0000398). This corresponded to the results of the analysis that was performed using STRING software in the *DiD+ exclusive* group of proteins, which computed a functional protein-protein interaction net (Fig. 7D; PPI enrichment *p* < 1.0e-16). There were two major hubs in the network that contained proteins that are involved in mRNA processing (GO: 0006397) and translation (GO: 0006412) or the mRNA catabolic process (GO: 0006402). These results suggest that stroma-derived cellular vesicles that were transferred through TNTs to CML cells can serve as a source of proteins to modulate gene expression at the level of mRNA processing and translation in CML cells.

To investigate the putatively modulated biological processes more comprehensively, the whole groups of proteins that were identified in the DiD+ and DiD- samples underwent a statistical overrepresentation test. We then combined the lists of results of the GeneOntology biological processes and found 41 biological processes for which the proteins were enriched in both samples with similar fold enrichment values (Supplementary Table S4), 16 biological processes that were enriched exclusively in the DiD- sample (Supplementary Table S4) and 15 biological processes that were enriched exclusively in the DiD+ sample (Supplementary Table S4, Fig. 7E). The latter ones were considered to be specifically modulated by proteins that were transferred in TNT-trafficked cellular vesicles. Again, the processes that are involved in the regulation of gene expression at the level of mRNA processing and translation were strongly represented (e.g., the regulation of mRNA splicing *via* spliceosome [GO: 0048024] and regulation of translation [GO: 0006417]). Additionally, new processes were identified: regulation of apoptotic signaling pathway (GO: 1902175 and 2001243), cell redox homeostasis (GO: 0045454), and response to drug (GO: 0042493). The enrichment of these processes might specifically promote survival and the response to conditions of stress in leukemic cells.

Altogether, our data showed that stromal cells conferred protection against the imatinib-induced apoptosis of CML cells by direct cell-to-cell trafficking of cellular vesicles and specific sets of proteins that can support cytoprotective and anti-apoptotic responses; with TNTs being involved in the observed exchange. These proteins can either directly help CML cells cope with the response to imatinib (through such processes as regulation of the apoptotic signaling pathway, cell redox homeostasis, and response to drug) or indirectly remodel their proteome to become enriched in proteins that are needed for the adaptation and response to imatinib. The latter possibility is attributable to the transfer of proteins that are involved in the regulation of gene expression at the levels of both transcription and translation. Our data suggest a novel mechanism of leukemia-stroma cross-talk by proteome exchange and may contribute to the development of new approaches to investigate the influence of the bone marrow stroma microenvironment on the leukemia response to treatment.

## Discussion

Cytoprotective signaling that is provided by the bone marrow niche toward leukemic cells has been well documented in different studies (Weisberg *et al*, 2008; Kumar *et al*, 2017b; Xu *et al*, 2016; Kumar *et al*, 2017a; Zhang *et al*, 2013). Considering that components of the leukemia microenvironment have been proposed as therapeutic targets (Krause *et al*, 2013; Duarte *et al*, 2018; Weisberg *et al*, 2008; Kumar *et al*, 2017b, 2017a; Zhang *et al*, 2013), understanding and characterizing interactions between different cell types in the leukemia-transformed bone marrow niche are critical for the development of novel therapeutic strategies. The present study provides new insights into leukemia-stroma crosstalk and highlights the important role of a novel, recently discovered mechanism of direct cell-cell communication *via* TNTs (Rustom *et al*, 2004; Önfelt *et al*, 2004; Eugenin *et al*, 2009; Gousset *et al*, 2009). The present data showed that TNTs are formed between stromal cells and leukemic cells and actively participate in the bidirectional transfer of vesicles and proteins. The transfer from stromal cells to leukemic cells resulted in cytoprotection and resistance to imatinib, thus making our data clinically relevant.

We found evidence that membrane connections that are formed between leukemic cells and stromal cells possess all features that are necessary to be classified as TNTs. To date, no specific molecular markers of TNTs have been developed. Thus, the presence of several characteristic morphological features is required to properly identify intercellular connections as TNT structures (Austefjord *et al*, 2014; Ariazi *et al*, 2017). Tunneling nanotubes that were found in the present study linked two distant cells and contained F-actin. Some of the TNTs also possessed microtubules and molecular motors (e.g., myosins). Moreover, membranes and actin filaments from both stromal cells and leukemic cells contributed to TNT formation. Together with our time-lapse recordings, these results support one of the mechanisms that were proposed for TNT formation based on “cell dislodgement” (Onfelt *et al*, 2006; Davis & Sowinski, 2008). We found that two cells first closely apposed each other, followed by the formation of a nanotube while they moved apart. Similar observations have been reported for immune cells (Önfelt *et al*, 2004; Onfelt *et al*, 2006; Watkins & Salter, 2005) and cells of other origin (Reichert *et al*, 2016). Because of their similarities, the morphology of TNTs has often been compared to filopodia, but their differences and unique features are quite clear (Delage *et al*, 2016).

Based on 3D reconstructions of confocal images, we excluded the possibility that the structures that formed between leukemic cells and stromal cells are filopodia-like protrusions, based on the findings that they did not adhere to the substratum, provided plasma membrane continuity between cells, and facilitated cargo transfer. Additionally, the lengths and diameters of TNTs remained within ranges that have been previously reported (Austefjord *et al*, 2014; Sisakhtnezhad & Khosravi, 2015). Still more advanced studies are needed to verify whether nanotubes formed between stromal and leukemic cells all possess open ends, form a synaptic nanotubular connections allowing for cargo transfer or represent a mixture of both types. This is still an on-going debate in the field. However, lack of such final conclusion in this term does not interfere with our findings and the possible role in the leukemia-stroma cross-talk.

We found that co-cultures of leukemic cells and stromal cells increased the number and activity of TNTs that formed between stromal cells and leukemic cells. Interestingly, leukemic cells are unable to efficiently form TNTs between each other. However, the presence of stromal cells stimulated leukemic cells to participate in TNT formation, including the involvement of their plasma membrane and cytoskeletal components. Even if the precise mechanism of TNT formation between leukemic cells and stromal cells is still unclear, we found strong evidence that the percentage of leukemic cells that are involved in TNT formation and the percentage of leukemic cells that received cargo increased under co-culture conditions. Our hypothesis that the ability of leukemic cells to form TNTs is stimulated by the microenvironment is supported by previous studies, in which AML cells had a low number of TNTs in general, but once they were isolated from the bone marrow of patients, a higher number of TNT/100 cells was found. This was not observed in AML cells that were cultured without any stromal component (Omsland *et al*, 2017). In AML cells that were cultured alone, the number of TNT/100 cells was very low and similar to the TNT index that was observed by us in mono-cultures. However, the authors of this previous study did not investigate leukemia-stroma interactions. In another study of leukemia, only cargo transfer toward mesenchymal cells was investigated, and the number of TNTs per cell was not assessed (Polak *et al*, 2015).

Furthermore, in the present study, the nanotubes that formed between leukemic cells and stromal cells mediated the transfer of different types of cargo, such as mitochondria and vesicles. Using different approaches, including fluorescence confocal microscopy with 3D reconstruction, and correlative light-electron microscopy with electron tomography image reconstruction, we found vesicles inside the lumen of TNT and confirmed their existence inside TNT by CLEM electron tomography. Their transfer between cells was confirmed by time-lapse micrscopy. Membrane vesicles have been previously shown to be nanotube cargo (Rustom *et al*, 2004; Gurke *et al*, 2008). The TNT-mediated transfer of vesicles that was observed in the present study has been previously reported (Rustom *et al*, 2004; Gurke *et al*, 2008; Delage *et al*, 2016), however without any evidence about the biological function of such transfer. We showed that the vesicles transferred from stromal to leukemic cells mediate resistance to imatinib. Moreover, the velocity of vesicles that transferred within TNTs was similar to previous studies (Gurke *et al*, 2008; Bénard *et al*, 2015). The commonly used methodology that is based on the fluorescent tracking of vesicles using DiD dye in donor cells combined with the flow cytometry analysis of DiD fluorescence in acceptor recipients (Gurke *et al*, 2008; Gousset *et al*, 2013; Abounit *et al*, 2015) has been used by us to quantify vesicle exchange, upon exclusion of indirect extracellular vesicles transfer. We found that the bidirectional transfer of vesicles between leukemic cells and stromal cells was not equal in both directions and strictly regulated, especially in heterotypic cellular networks. Importantly, we excluded the possibility of vesicle exchange through the secretion of vesicles using two commonly used control setups: the transwell system (in which two types of cells were physically separated) and the treatment of acceptor cells in conditioned medium that was collected from donor cells that contained all types of secreted vesicles (Abounit *et al*, 2016a). Thus, we confirmed that physical, direct cell-to-cell contact is necessary for the efficient transfer of vesicles. As no general pharmacological inhibitors of TNT activity have been discovered to date, such controls are necessary to verify whether the observed cargo transfer is TNT-mediated.

One of the most interesting findings in the present study was that the TNT-mediated transfer of vesicles from stromal cells to leukemic cells correlated with the protection of leukemic acceptor cells from imatinib-induced cell death. Based on these data we can hypothesize that the cell-to-cell TNT-mediated transfer of vesicles has biological significance and influences resistance to imatinib as part of the stroma-provided protection. Exchange of vesicles has been already shown, but mostly as a model cargo to study the efficiency and mechanism of TNT formation and function. Our data clearly showed that the cell-to-cell-mediated transfer of vesicles might be a novel element of the stroma-provided cytoprotection of leukemic cells. Additionally, mitochondrial transfer within the bone marrow microenvironment was summarized in a recent review by Griessinger, who proposed that receiving mitochondria by damaged cells from healthy donors may be a potent mechanism of survival, cytoprotection, and regeneration (Griessinger *et al*, 2017). The possibility of the TNT-mediated transfer of vesicular cargo between leukemic cells and stromal cells is supported by the finding that signaling from acute lymphoblastic leukemic cells to primary mesenchymal stem cells in the bone marrow microenvironment occurs through TNTs. This signaling led to an increase in the secretion of proleukemic and prosurvival cytokines by stromal cells. Still unclear, however, is the way in which such signaling can influence the secretion of soluble factors (Polak *et al*, 2015).

Our trans-SILAC experiments, combined with flow cytometry and fluorescent sorting, showed that functional sets of proteins can be transferred directly together with vesicles from stromal cells to leukemic cells. The transfer of single proteins via TNTs has been previously reported (Abounit *et al*, 2016a, 2016b). We further performed a proteomic analysis of transferred proteins, demonstrating that whole sets of proteins can be exchanged. The finding that such proteins are not transferred randomly but are likely specifically selected for transport is novel and might have significant biological implications. The bioinformatic analysis of transferred proteins indicated that they can form functional protein-protein association networks and regulate specific biological processes that are important for cellular adaptation and survival. This may be meaningful in the context of cancer treatment and the response to chemotherapy. The transfer of proteins from stromal cells to leukemic cells may significantly support the resistance to treatment and contribute to disease progression. Such protein transfer to increase cytoprotection, in addition to the transfer of mitochondria from healthy cells to damaged cells in monotypic and heterotypic biological setups, likely increases the survival of recipient cells (Li *et al*, 2017; Sanchez *et al*, 2017; Wang & Gerdes, 2015; Pasquier *et al*, 2013). Altogether, our data confirmed the general role of direct intercellular cargo transfer as a supportive and protective mechanism.

Our findings support the hypothesis of intercellular proteome exchange. This hypothesis postulates that the proteome of autonomic cells does not exist under physiological conditions because no cell is a separate “island” (Rechavi *et al*, 2009; Ahmed & Xiang, 2011). Our data clearly showed that proteomic interactions should be considered in studies of intercellular interactions and cellular networks.

In summary, we unveiled a novel role for TNTs in the protection of leukemic cells by stroma. Direct cell-contact-dependent vesicle transfer correlated with resistance to the therapeutic drug imatinib. Together with the transfer of specific sets of proteins that play roles in cellular adaptation and survival, we might propose that TNT signaling plays a significant cytoprotective role. This observation is relevant with regard to the resistance of leukemic cells to imatinib, including leukemia stem cells that reside in bone marrow (Lane *et al*, 2009). Various protective mechanisms that participate in this phenomenon have been proposed. Tunneling nanotube-mediated cross-talk may be very potent mechanism that participates in the protection of leukemic cells and if so, then TNT-mediated communication would be a potential therapeutic target for the treatment of leukemia.

## Materials and Methods

### Cell lines and reagents

K562 and HS-5 cell lines were obtained from the American Type Culture Collection and cultured in RPMI medium that was supplemented with 10% fetal bovine serum (FBS), 1% L-glutamine, and 1% penicillin streptomycin. The cells underwent a regular screen for *Mycoplasma* contamination (i.e., the polymerase chain reaction-based detection of *Mycoplasma*). The K562 GFP cell line was established by Dr. M. Kusio-Kobiałka. Imatinib was a generous gift from the Pharmaceutical Research Institute (Warsaw) and used at concentrations of 0.5, 1, and 5 µM.

### Co-culture system and flow cytometry measurements

#### Exchange of cargo between cells

Donor cells were labelled with DiD (ThermoFisher Scientific; 1.5 μl/1 ml of cell culture medium) for 15 min at 37°C, washed twice in PBS, and plated in fresh cell culture medium for an additional 16 h. Afterward, the cells were harvested, counted, and seeded in co-culture with acceptor cells in 12-well cell culture plates (1 × 10^5^ HS-5 cells plus 0.8 × 10^5^ K562 GFP cells) to reach a 1:1 ratio after 24 h. For flow cytometry, all of the cells were harvested and subjected to analysis using a BD LSRFortessa cytometer. The flow cytometry data were further analyzed using Diva and FlowJo software.

#### Trans-well and CM controls

To physically separate donor and acceptor cells in co-culture, HS-5 and K562 cells were plated in the lower and upper chambers of a transwell system (ThinCert, Greiner Bio-One, 1 µm pores, 2 × 10^6^ pores/cm^2^) that were separated by a filter with 1 μm pores. After 24 h of co-culture, the cells from both chambers were harvested and underwent flow cytometry analysis together with cells that were co-cultured without physical separation.

As a control for the conditioned media, donor cells were labeled as described above and plated for 24 h at the same density as in the exchange experiment. After 24 h of cell culture, the supernatant was collected, centrifuged to remove cells and cellular debris, and added to acceptor cells in 12-well culture plates. After another 24 h, acceptor cells were harvested and underwent flow cytometry analysis.

#### Flow cytometry analysis of K562 acceptor cells

To investigate the apoptosis of K562 acceptor cells, co-cultures of DiD-labeled HS-5 cells with K562 GFP cells were untreated or treated with imatinib for 48 h and stained with AnnexinV-PE and 7-AAD (BD Pharmingen) according to the manufacturer’s instructions and analyzed using BD LSRFortessa. Acceptor cells that were positive and negative for transported cargo were separated by gating and separately analyzed for apoptosis. To study caspase activation, cells from co-cultures were labeled with Violet Live Cell Caspase Probe (BD Pharmingen) according to the manufacturer’s instructions and 7-AAD for live cell discrimination. Acceptor cells that were positive and negative for transported cargo were separated by gating, and the percentage of cells with active caspases was calculated. To investigate proliferation, K562 acceptor cells were labeled with 10 μM Cell Proliferation Dye eFluor 450 (eBioscience) and analyzed after 24, 48, and 72 h of co-culture with DiD-stained HS-5 donor cells. In analogous experiments, in which DiD-labeled HS-5 cells were donors and Proliferation Dye-stained K562 cells were acceptors, mitochondrial mass, reactive oxygen species production, and mitochondrial membrane potential were investigated using MitoTracker Green (100 nM for 30 min at 37°C and 5% CO_2_), H_2_DCF-DA (2 μM for 15 min at 37°C and 5% CO_2_), and JC-1 (2 μM for 10 min at 37°C and 5% CO_2_), respectively.

### Fluorescent imaging and live cell microscopy

#### Immunocytochemistry and immunofluorescence

Cells were plated for 24-48 h on poly-L-lysine-coated coverslips, fixed with 4% paraformaldehyde (PFA) that was added directly to the cell culture medium, permeabilized with 0.1% Triton X-100 for 3 min, blocked with 5% FBS for 1 h, and incubated with appropriate antibodies and fluorescent stains. Phalloidin was used for actin staining, and DAPI was used for nuclear labeling. Microtubules were labeled with monoclonal anti-β-tubulin antibody (Sigma-Aldrich), MyoVa antibody, (Cell Signaling Technology), MyoVI antibody (Proteus), and MyoVIIa antibody (Proteus). Mitochondria and cellular vesicles were labeled the day before cell plating with 250 nM MitoTracker Deep Red or DiD, respectively (both from ThermoFisher Scientific). Images were acquired using a Zeiss LSM 780 microscope with a 63× objective and further processed using ImageJ and Imaris software.

#### Tunneling nanotube imaging in living cells

Cells were plated on Lab-Tek Chamber Slides (ThermoFisher Scientific) that were coated with poly-L-lysine. To distinguish K562 cells from HS-5 cells, the K562 GFP cell line was used. Plasma membranes were labeled with Wheat Germ Agglutinin, and Alexa Fluor 647 Conjugate (ThermoFisher Scientific) was added directly to the chamber 10 min before imaging. Images were acquired using an SP8 Leica microscope with a 63× or 100× objective.

#### Tunneling nanotube quantification

Cells were plated and labeled as described above. Ten fields of view (155 μm × 155 μm) with z-stacks that covered the majority of the cell volume were acquired using an SP8 Leica microscope with a 63× or 100× objective. The data were manually analyzed using ImageJ software with the Cell Counter plugin.

### Plasma membrane contribution and actin penetration experiments

To assess the contribution of the plasma membrane of CML cells and actin penetration into TNTs, K562 cells were transfected by nucleofection (Amaxa Nucleofector Technology, Lonza) using a GPI-GFP plasmid (kind gift from D. Davis) or LifeAct-GFP plasmid (kind gift from J. Włodarczyk), respectively. Twenty-four hours after transfection, the cells were sorted based on GFP fluorescence and co-cultured with HS-5 cells on poly-L-lysine-coated Lab-Tek Chamber Slides (ThermoFisher Scientific) for 48 h. Images were acquired using an SP8 Leica microscope with a 63× objective and analyzed using ImageJ software.

### Electron microscopy

#### Scanning electron microscopy

Cells were plated on poly-L-lysine-coated coverslips. After 48 h, the cells were fixed with 2.5% glutaraldehyde and 2% PFA in PBS for 20 min at room temperature. The cells were then washed twice with PBS and then twice with water and dehydrated by subsequent washes in an ascending series of ethanol concentrations (50%, 70%, 96%, and 99.9%, 10 min each). The samples were then subjected to critical-point drying, gold-coated, and imaged on 3View using Zeiss Sigma VP SEM column. The secondary electron signal was used to obtain an image.

#### Transmission electron microscopy

Monolayers of cells were seeded on IBIDI-gridded, glass window, poly-L-lysine-coated plates. After 24 h, the cells were labeled with WGA AF-647 and fixed in 2.5% glutaraldehyde and 2% PFA in cacodylate buffer. Light microscopy images were then acquired using an Olympus IX81 widefield microscope using the software CellR. The cells were then washed with cacodylate, contrasted in 1% osmium and 1% uranyl acetate (UA), dehydrated in ethanol, and embedded in Epon resin. The fixation, contrasting, and dehydration steps were performed in a PELCO Biowave Pro microwave (Lorentzen *et al*, 2018). Next, 50 serial sections (300 nm thick) were collected on formvar-coated slot grids, 15 nm gold beads were added as fiducial for tomogram reconstruction, and then sections were poststained in 2% UA and lead citrate. Regions of interest were imaged using a FEI Tecnai F30 (tomography) electron microscope, operated at 300 kV. Image were acquired with a OneView Camara (Gatan). Image montages and tomogram reconstruction processing were performed using the IMOD software package (Kremer *et al*, 1996).

### Trans-SILAC

#### Cell labeling with heavy isotopologues of lysine and arginine

The SILAC medium was supplemented with 10% of dialyzed FBS, 1% Pen/Strep, 0.274 mM L-lysine, and 1.15 mM L-arginine and filtered (0.22 µm pores). Donor cells were labeled with heavy isotopologues of lysine and arginine: L-lysine:2HCL (13C6, 99%; 15N2, 99%) and L-arginine:HCL (13C6, 99%; 15N4, 99%; Cambridge Isotope Laboratories). Cells were maintained in the appropriate medium for 9 days to enable complete labeling of the proteome. The medium was changed every 2-3 days. On day 8, donor cells were labeled with DiD as described above. On day 9, co-cultures that were untreated or treated with 1 μM imatinib were established. This experiment was performed in duplicate.

#### Co-cultures and cell sorting

Twenty-four hour co-cultures were sorted using a BD FACSAria sorter. Green fluorescent protein fluorescence was excited with a 488 nm laser, and emission was detected with a 530/30 filter. DiD fluorescence was excited with a 633 nm laser, and emission was detected with a 660/20 filter. Donor and acceptor cells were separated by FACS, and acceptor cells were further sorted into two subpopulations: positive and negative for transferred cargo. Cell pellets were immediately frozen in liquid nitrogen and stored at -80°C until further sample preparation.

#### Sample preparation

Cell pellets were lysed in 25 mM ammonium carbonate and 0.1% Rapigest (pH 8.2). Protein lysate (10 μg) was digested with Trypsin Gold, reduced with 5 mM tris(2-carboxyethyl)phosphine (TCEP), and blocked with 5 mM iodoacetamide. For a more in-depth analysis, DID+ and DID- samples were diluted with 4× Laemmli buffer, boiled for 5 min at 95°C, and loaded onto a sodium dodecyl sulfate-polyacrylamide gel electrophoresis gel. Lanes were cut into six pieces, and proteins were digested using trypsin according to standard protocols (Shevchenko *et al*, 2007). Peptides were extracted, purified by styrenedivinylbenzene reverse-phase sulfonate (SDB-RP); also known as mixed mode chromatography stage tips (Kulak *et al*, 2014), and stored at 4°C prior to MS analysis.

#### LC-MS/MS analysis

Mass spectrometry analysis was performed using a nanoAcquity UPLC system (Waters) that was directly coupled to a Q Exactive mass spectrometer (ThermoFisher Scientific). Peptides were separated by a 180-min linear gradient of 95% solution A (0.1% formic acid in water) to 35% solution B (acetonitrile and 0.1% formic acid). The measurement of each sample was preceded by three washing runs to avoid cross-contamination. The mass spectrometer was operated in the data-dependent MS-MS2 mode. Data were acquired in the *m*/*z* range of 300-2000 or 300-1750 at a nominal resolution of 70,000.

Data were analyzed using the Max-Quant 1.5.3.12 platform. The reference human proteome database from UniProt was used. False discovery rates of protein and peptide-spectrum matches (PSM) levels were estimated using the target-decoy approach at 0.01% (protein FDR) and 0.01% (PSM FDR), respectively. The minimal peptide length was set to 7 amino acids, and carbamidomethyolation at cysteine residues was considered a fixed modification. Oxidation (M) and Acetyl (Protein N-term) were included as variable modifications. Only proteins that were represented by at least two unique peptides in two biological replicates are shown and were further considered. The data analysis was performed using MaxQuant software and the MaxLFQ algorithm.

#### Analysis of LC-MS/MS data

Only proteins that were identified in both biological replicates underwent further analysis. Lists of proteins were analyzed using the Panther application for GeneOntology software (Mi *et al*, 2017), STRING-confidence view (Szklarczyk *et al*, 2015), and Venny 2.1. Venny is an interactive tool that is used to compare lists with Venn diagrams (http://bioinfogp.cnb.csic.es/tools/venny/index.html). Additionally, lists of proteins were grouped according to their molecular weights based on the UniProt database. The mass spectrometry data from this publication have been deposited to the ProteomeXchange Consortium via the PRIDE [https://www.ebi.ac.uk/pride] partner repository with the dataset identifier PXD011013.

### Statistical analysis

All of the experiments were performed in at least three independent repetitions. All of the data are presented as mean ± SEM. Student’s *t*-test with Welch correction was used to test differences between two conditions. Values of *p* < 0.05 were considered statistically significant. Microsoft Excel and GraphPad Prism software were used for the data analysis.

## Acknowledgements

We thank Piotr Sunderland for initial help with live cell imaging using the SP8 Leica microscope. The microscopy experiments were performed at the Laboratory of Imaging Tissue Structure and Functions at the Nencki Institute of Experimental Biology in Warsaw and the D. Davis laboratory at the University of Manchester. Transmission electron microscopy was performed at the Laboratory of Electron Microscopy at the Nencki Institute in Warsaw; CLEM, and electron tomography analyses were performed at the EMBL Electron Microscopy Core Facility in Heidelberg. The mass spectrometry analysis was performed at the Laboratory of Mass Spectrometry, Institute of Biochemistry and Biophysics, Polish Academy of Sciences, in Warsaw and at the CECAD Research Center at the University of Cologne. This study was supported by grants from the National Science Center, Poland (no. 2013/10/E/NZ3/00673 to KP, 2015/17/B/NZ3/00557 to JW, and 2017/25/N/NZ3/00741 to MDK), and the Manchester Collaborative Centre for Inflammation Research (funded by a precompetitive open-innovation award from GlaxoSmithKline, AstraZeneca, and the University of Manchester).

## Author contributions

KP coordinated and designed the studies, discussed the data, and wrote the manuscript. MDK designed and performed most of the experiments, analyzed the data, and wrote the manuscript. WD performed the CLEM experiments under supervision of PR and YS; WD discussed the data and together with MDK performed SEM experiments. PR and YS were involved in the CLEM experiments, discussed the data and edited the manuscript. JW discussed the mass spectrometry experiments and supervised MZ-K. MZ-K prepared the samples for MS and analyzed the MS data. LT performed the caspase activity assay under supervision of MDK. AK was responsible for cell sorting and participated in flow cytometry analyses. KS was involved in microscopy experiments and imaging of TNTs together with MDK. DMD supervised KS and was involved in data discussion. All of the authors contributed to editing the manuscript.

## Supplementary Figure Legends

**Supplementary Figure S1.** Diameters and lengths of homo- and heterotypic TNTs that were measured either in living cells by confocal microscopy (**A, B**) or in fixed SEM samples (**C, D**). Homotypic TNT formed by stromal cells (HH) or leukemic cells (KK) as well as heterotypic TNTs (HK) were measured.

**Supplementary Figure S2.** Average lengths of heterotypic TNTs that depended on the origin of the plasma membrane that constituted a given TNT.

**Supplementary Figure S3.** Percentage of apoptotic leukemic cells upon treatment with imatinib either cultured alone or in co-culture with stromal cells.

**Supplementary Figure S4.** The Scheme showing a mass spectrometry (MS)-based trans-SILAC approach performed to identify proteins that are shuttled between CML cells and stromal cells.

**Supplementary Figure S5. A.** Confidence view of functional protein association networks (String software) of proteins that were transferred toward stromal cells from CML cells in a co-culture set-up. Hubs represent proteins. Edges represent protein-protein associations. The line thickness indicates the strength of data support. PPI enrichment *p* = 3.6e-08. **B.** Confidence view of functional protein association networks (String software) of proteins that were transferred toward CML cells from stromal cells in a co-culture set-up. PPI enrichment *p* < 1.0e-16. **C.** Results of statistical overrepresentation test of the list of proteins that were transferred toward stromal cells from CML cells (Panther GeneOntology). The list was tested for the enrichment of proteins that are involved in GeneOntology Biological Processes against the reference list of all *Homo sapiens* genes in the database. Fisher’s Exact Test with FDR multiple test correction was used. Only results with FDR ≤ 0.05 are presented. **D.** Results of statistical overrepresentation test on the list of proteins that were transferred toward CML cells from stromal cells (Panther GeneOntology). The list was tested for the enrichment of proteins that are involved in GeneOntology Biological Processes against the reference list of genes of the K562 cell line from the PRIDE database, completed with the list of genes that were identified in the present study. Fisher’s Exact Test with FDR multiple test correction was used. Only results with FDR ≤ 0.05 are presented.

